# Additive Manufacturing Leveraged Microfluidic Setup for Sample to Answer Colorimetric Detection of Pathogens

**DOI:** 10.1101/2023.08.10.552726

**Authors:** Sripadh Guptha Yedire, Imman Isaac Hosseini, Hamed Shieh, Arash Khorrami Jahromi, Tamer AbdelFatah, Mahsa Jalali, Sara Mahshid

## Abstract

Colorimetric readout for the detection of infectious diseases is gaining traction at the point of care/need owing to its ease of analysis and interpretation, and integration potential with highly specific Loop-mediated amplification (LAMP) assays. However, coupling colorimetric readout with LAMP is rife with challenges including, rapidity, inter-user variability, colorimetric signal quantification, and user involvement in sequential steps of the LAMP assay, hindering its application. To address these challenges, for the first time, we propose a remotely smartphone-operated automated setup consisting of (i) an additively manufactured microfluidic cartridge, (ii) a portable reflected-light imaging setup with controlled epi-illumination (PRICE) module, and (iii) a control and data analysis module. The microfluidic cartridge facilitates sample collection, lysis, mixing of amplification reagents stored on-chip, and subsequent isothermal heating for initiation of amplification in a novel way by employing tunable elastomeric chambers and auxiliary components (heaters and linear actuators). PRICE offers a new imaging setup that captures the colorimetric change of the amplification media over a plasmonic nanostructured substrate in a controlled and noise-free environment for rapid minute-scale nucleic acid detection. The control and data analysis module employs microprocessors to automate cartridge operation in tandem with the imaging module. The different device components were characterized individually and finally, as a proof of concept, SARS-CoV-2 wild-type RNA was detected with a turnaround time of 13 minutes, showing the device’s clinical feasibility. The suggested automated device can be adopted in future iterations for other detection and molecular assays that require sequential fluid handling steps.

## 1. Introduction

The colorimetric signal transduction technique has been extensively applied for the detection of biological analytes.^1,2^ Colorimetric readout techniques are increasingly adopted for pathogen detection at the point of care (POC) ^3–5^, owing to key features like ease of analysis and interpretation, binary end-point detection, and affordability.^6–10^ In the context of pathogen detection, colorimetric readout coupled nucleic acid amplification assays are gaining traction.^11,12^ Specifically, colorimetric loop-mediated isothermal amplification (LAMP) is increasingly adopted ^13,14^, due to a constant temperature for amplification^15^, PCR comparable specificity and sensitivity^16^, and stability against some amplification inhibitors.^17^ However, the POC deployment of colorimetric readout coupled LAMP is rife with challenges like inter-user variability, signal quantification, and the requirement of trained personnel in assay deployment.

To offset the drawbacks of inter-user variability and signal quantification, there is increasing adoption of smartphones and open-source technologies^9,18–25^, enabling miniaturization and ease of data collection and analysis.^26,27^ However, smartphones have rigidity in imaging parameters and are prone to interference from changes in ambient light^28^, ultimately affecting the data collection and analysis. Additionally, inherent restrictions in smartphones for working in color spaces like RGB, HSV, CIELAB, and accessibility to raw data might hamper signal quantification.^29^ Further, previously reported imaging setups lack control over spatial and intensity characteristics of the illumination for image capture.

One decisive feature of a POC test is integration of sequential steps of nucleic acid amplification i.e., sample collection, sample processing, reagent mixing, amplification reaction, and detection sub-steps.^30^ To address these broad needs, microfluidic setups could help implement techniques that allow metering and precise control of fluids as well as effective heat transfer, onto a single platform.^31–33^ Moreover, confined reaction volumes allow for fast and high throughput analysis, and enhanced heat transfer.^32,34,35^ Although several microfluidic amplification-based platforms were reported, they often involve user involvement in one or more of these sub-steps, especially in sample collection, sample preprocessing, and/or fluid manipulation steps.^36–40^ An exciting subset of fluid manipulation techniques based on finger/hand-powered actuation gained traction in recent years for a wide range of biomedical applications.^41–43^ These techniques rely on the generation of pressure gradient in the microchannels via a manual operation by the user via deformable elastomeric chamber^44,45^ or solid piston actuation^46^, leading to pumping in the microchannels. It is worth noting, these techniques may address some challenges such as complex design strategies, variability among users, the requirement of an external operator, and complicated micro-valving setups often require extensive microfabrication and assembly processes.^47–49^

Additive manufacturing (3D printing) techniques can substantially address the challenges associated with microfabrication and with microfluidic handling on a large scale.^50,51^ Indeed AM can bring about outcomes comparable to traditional lithography in terms of temporal and spatial resolution, material, and mechanical properties while being cost-effective, rapid, and scalable.^52,53^ While the integration of LAMP onto additively manufactured devices is gaining traction^20,54–56^, some challenges still exist with the combination of all the sequential steps onto a modular user-friendly platform.

In this work, a smartphone-operated automated setup is employed to automate a colorimetric LAMP assay, from sample collection to data analysis. The setup is comprised of three main modules, (i) additively manufactured microfluidic cartridge, (ii) portable reflected-light imaging setup with controlled epi-illumination (PRICE) module, (iii) control and data analysis module. The microfluidic cartridge facilitates sample collection, lysis, mixing of amplification reagents stored on-chip, and subsequent isothermal heating for initiation of amplification. The fluid flow in the cartridge is mediated by angle dependent mechanical actuation of elastomeric chambers for suction-driven flow manipulation. The colorimetric change of the amplification media is captured with the PRICE module that facilitates imaging over a plasmonic nanostructured platform we previously reported for rapid minute-scale nucleic acid detection.^57^ The control and data analysis module comprising of Arduino and Raspberry Pi enables signal transduction via a concerted operation between PRICE, microfluidic cartridge, and auxiliary components (heaters and linear actuators). The different modules were characterized individually and finally, as a proof of concept, SARS-CoV-2 wild-type RNA was detected with a turnaround time of 13 minutes.

## 2. Experimental

### 2.1. Optical train setup

The portable reflected-light imaging setup with controlled epi-illumination (PRICE) has two main important submodules, (i) illumination submodule and (ii) image capture submodule. In the illumination submodule, a 5000 K 90CRI LUXEON LED (Lumileds Inc.) was used as the primary illumination source. To collimate the LED, a diffusive aspheric condenser lens (d= 25.4 mm, f=20.1 mm, Thorlabs) was used. A ring-actuated iris diaphragm (Thorlabs) was used as a field diaphragm with aperture diameters ranging from 8mm (minimum aperture opening) to 12 mm (maximum aperture opening). The collimated light is then illuminated on the back of an achromatic doublet lens (d= 25.4 mm, f= 30 mm, Thorlabs), creating an image at the focal length of the lens. A ring-actuated iris diaphragm (Thorlabs) was used as an aperture diaphragm with aperture diameters ranging from 8mm (minimum aperture opening) to 12mm (maximum aperture opening). Finally, the image of the light at the aperture diaphragm is collimated by an achromatic doublet lens (d= 25.4 mm, f= 30 mm, Thorlabs). The collimated light is then projected onto the back aperture of the objective (TU Plan Fluor EPI 20x, N.A. 0.45, W.D. 4.5 mm, Nikon Inc.) via a beam splitter (Reflectance: Transmittance-30:70, d= 25.4 mm, Thorlabs) placed at an angle of 45degrees with the vertical. The reflected light from the sample placed at the working distance is collected by the objective. Since the objective is infinity focused, a condenser lens (tube lens, f=200 mm) was used to project the image onto the CMOS sensor (Sony IMX477R, 12.3MP, Raspberry Pi Inc.).

### 2.2. Fabrication of the microfluidic cartridge

The microfluidic cartridge enables the integration of sample collection, sample lysis, reagent mixing and amplification steps on a single platform. All the microfluidic chip and the cartridge components were designed using AutoCAD™ and SolidWorks software. The first step of the fabrication process is patterning the fluidic channels (400 µm width and 50 µm thick) using UV photolithography. A lithography mask was designed to pattern a 50 µm thick SU-8 layer (SU-8 2050, MicroChem Corp., MA, USA) (Fig. S1a) on a silicon substrate (Fig. S1a) via a straightforward lithography process. In the second step (Fig. S1b), the QolorEX nanostructured platform^57^ (Fig. S1b (3)) was fabricated using a fabless nano-patterning technique. Initially, a generic method was employed to produce a colloidal self-assembly monolayer (SAM) of nanoparticles at a water/air interface. Following that, the honey-comb structures generated were moved to the Si Substrate. After that, a ZnO thin film (120 nm) was used to create a low dielectric constant back reflector. Finally, a thin layer of aluminum (10 nm) was applied to produce a tunable localized surface plasmon resonance on a white background. Fig. S1b (Inset) shows the various materials involved in the fabrication of the color-sensitive platform (Fig. S1b (3)). In the next step, a thin polydimethylsiloxane-PDMS layer (10:1, PDMS SYLGARD 184 silicone elastomer, Dow Consumer Solutions, QC, Canada) was punched with holes at channel inlets. The punched PDMS layer was plasma treated for 40sec at maximum power (Harrick Plasma cleaner, PDC-32G (115 V), 18 W). This surface activated PDMS layer was bonded to the SU-8 fluidic layer to create closed microfluidic channels (Fig. S1c (4)). Next, PDMS based elastomeric chambers layer was fabricated using SLA printed molds (Fig. S2). This elastomeric chamber layer was bonded to the fluidic-PDMS layer by plasma treatment for 40 sec at maximum power (Fig. S1d (5)). Finally, the Stereolithography (SLA) 3D printed cartridge (printed at 50 µm resolution with the Form 3, Formlabs, USA) is bonded to the PDMS covered microfluidic chip using a double-sided tape conducive to plasma activated bonding^58^[NO_PRINTED_FORM] (treated at maximum power for 40sec and placed in 95^0^ C for 90 min) (Fig. S1(e) (6&7) & Fig. 3.3(f)). The figure (Fig. S1(f)) shows the exploded view of the cartridge showing multiple components. The brass metal inserts (Fig. S1(f) (9&10)) are employed for lysis of the sample. Small screws (Fig. S1(f) (13)) are used to seal-off the inlets (in the printed cartridge) used for reagent loading. The components (Fig. S1 (f) (11&12)) are 3D printed screws for actuating the elastomeric chambers by applying a small lateral force during automation, resulting in negative pressure gradient and subsequent flow of liquid from the inlets.

All the 3D printed components are designed with SolidWorks software and fabricated using an SLA 3D printer (Form 3, Formlabs, USA) at a layer thickness of 25 µm in the z-axis as per the company’s specifications. The post-printing treatment included a wash in isopropyl alcohol (IPA) for 20 min followed by drying and UV curing at 60^0^ C for 20 min. Once the curing is done, the supports are removed. Following this, the collection funnel, lysis funnel, and reagent storage chambers housed in the 3D printed fluid handling attachment, are coated with Epoxy resin (Artresin, Canada) to make the surface biocompatible.^59^ Finally, a hollow biocompatible aluminum insert (McMaster-Carr Inc, Canada) is incorporated into the 3D printed fluid handling attachment as the lysis chamber.

### 2.3. Fabrication of elastomeric chambers

In this work we employed 3D printed molds for the fabrication of elastomeric chambers with PDMS. SolidWorks software was used to design the master mold. We employed high resolution SLA 3D printing (Form 3, Formlabs, USA) at a layer thickness of 25 µm in the z-axis as per the company’s specifications. The post-printing treatment included a wash in isopropyl alcohol (IPA) for 20 min followed by drying and UV curing at 60^0^ C for 20 min. Once the curing is done, the supports are removed. The master mold had two components, male and female molds (Fig. S2). Before proceeding to pour PDMS for molding, the 3D printed parts were surface treated by established protocol^60^ to avoid curing inhibition at the PDMS-mold interface.^61,62^ In brief, the 3D printed master mold post curing is first heated at 130^0^ C in oven for 4 h, following by an air plasma treatment (Harrick Plasma cleaner, PDC-32G (115 V), 18 W) for 3 min. The molds are then treated with 10 µl trichlorosilane (Sigma Aldrich) in a vacuum desiccator overnight. Finally, the PDMS-curing agent mixture in the ratio of 10:1 (PDMS to curing agent ratio), is poured into the female mold. The male mold fits tightly onto the female mold via the guides at the corners. The male mold has vertical openings to accommodate the overflow of excess PDMS upon insertion into the female mold (Fig. S2). The mold combination is then placed in the oven at 65^0^ C overnight for curing.

### 2.4. Electronic components of automation

The automation module had three main components, (i) Linear actuator system, (ii) Heating module, (iii) x-y translation stage and (iv) Imaging and data processing module (Fig. S3). All these components are controlled with, Arduino UNO (Arduino Inc.) and Raspberry Pi 4 (Raspberry Inc.). Five linear actuators (Actuonix Inc.) were employed to facilitate sequential fluid handling steps. The lysis and amplification were facilitated by a portable solder iron (TS-100) for saliva lysis and ceramic thermal heater (Bolsen Tech Inc.), respectively. The X-Y translation stage was realised using a linearly guided CNC stage driven by a stepper motor (FUYU Inc.), in combination with a continuous servo (SPT digital, SPT5325LV) contraption. The imaging and data processing is done on-board the Raspberry Pi 4. The CMOS sensor (Sony IMX477R, 12.3MP, Raspberry Pi Inc.), captures the images and is processed on board. The connection overview of electronics is depicted in Fig. S4. To realize an automated, user-friendly and wireless operation, a mobile application was created using a MIT app inventor, that allows the user to initialize, monitor and obtain the results (Fig. S5).

### 2.5. RT-LAMP assay

The RT-LAMP assay employed in this work uses the primer set against the ORF1ab gene obtained from Sigma-Aldrich, USA. The individual oligonucleotide concentrations of the 10x primer mix were, 0.2 µmol/L of forward outer primer, 1.6 µmol/L of Forward inner primer and backward inner primer, 0.4 µmol/L of forward loop primer and backward loop primer. The primer mix was mixed with WarmStart^®^ Colorimetric LAMP 2X Master Mix (NewEngland Biolabs, MA, USA), and RNase free water (Thermo Fischer Scientific, MA, USA). The standard reaction volume is 20 µl that consisted of 2 µl 10X primer mix, 10 µl 2X master mix, 7 µl RNase free water, and 1 µl synthetic SARS-CoV2 RNA (VR-3276SD ATCC, VA, USA) sample. The samples were incubated at 65°C for different periods to visualize color change versus time for different samples.

## 3. Results and discussion

### 3.1. Overview of operation

The current pathogen detection system has three major steps, (i) sample collection and system initiation, (ii) amplification assay, and (iii) image capture and data analysis (Fig. 1). In the first step, the user loads the analyte up to the specified level, and then close it with a tightly sealed lid. Following this, the user opens the sliding door of the benchtop automated setup and places the microfluidic cartridge on the stage holder. The user then connects to the automated setup via the developed mobile application and initiates the sample-to-answer workflow (i). A concerted operation between different components inside the box will facilitate all the key assay steps namely, sample lysis, mixing with amplification reagents, and heating for the amplification reaction (ii). Once the assay is completed, the colorimetric endpoint is imaged, and data is analyzed. The results are then transmitted to the user’s mobile application (iii).

**Fig 1.**
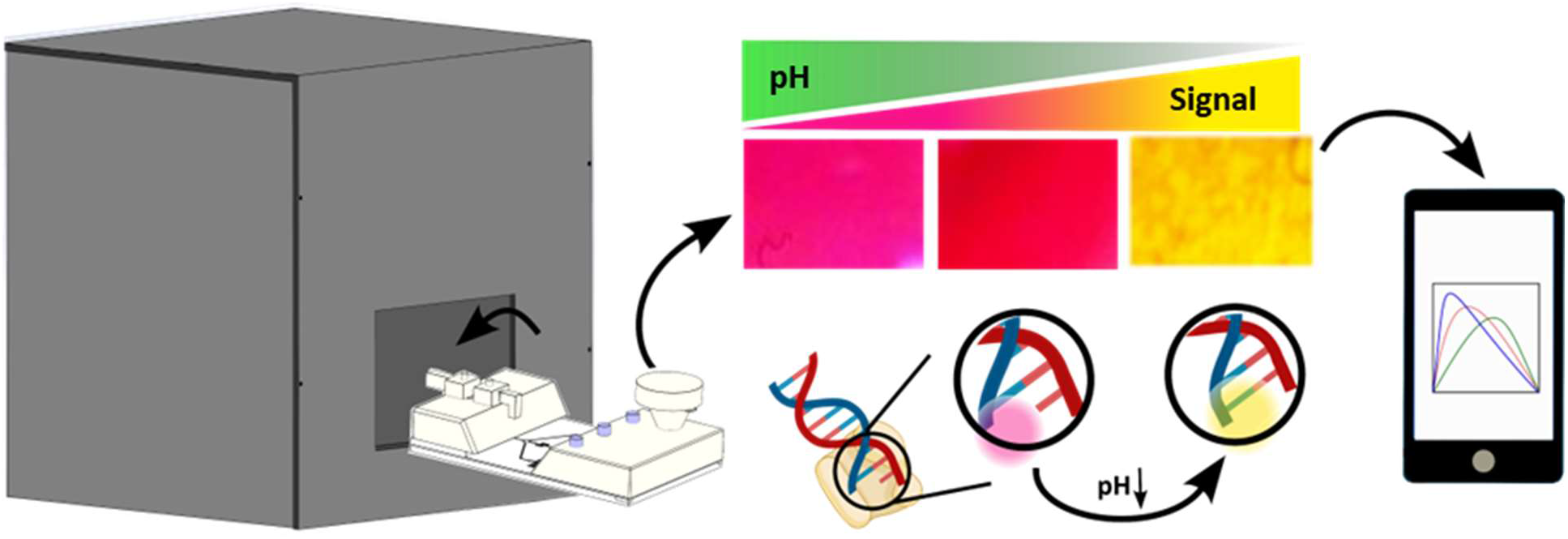
Schematic overview. The major steps of the automated workflow for pathogen detection

### 3.2. Development of the sample-to-answer workflow

The automated setup comprises three main modules, (i) imaging module (Fig. 2a(α)), (ii) microfluidic cartridge (Fig. 2a(β)), (iii) control and data analysis module (Fig. 2a(γ)), all housed in a benchtop and portable setup. The topmost module, the imaging module (Fig. 2a(α)), dubbed, portable reflected-light imaging setup with controlled epi-illumination (PRICE) was designed to capture the colorimetric change of the amplification reaction over a plasmonic nanostructured platform. The control and data analysis module γ further is comprised of δ-linear actuator unit and θ-microcontrollers and auxiliary electronic components (Fig. 2a). All these components-α, β, γ, θ, δ-work in tandem to enable a sample to answer workflow (Fig. 2b).

**Fig 2.**
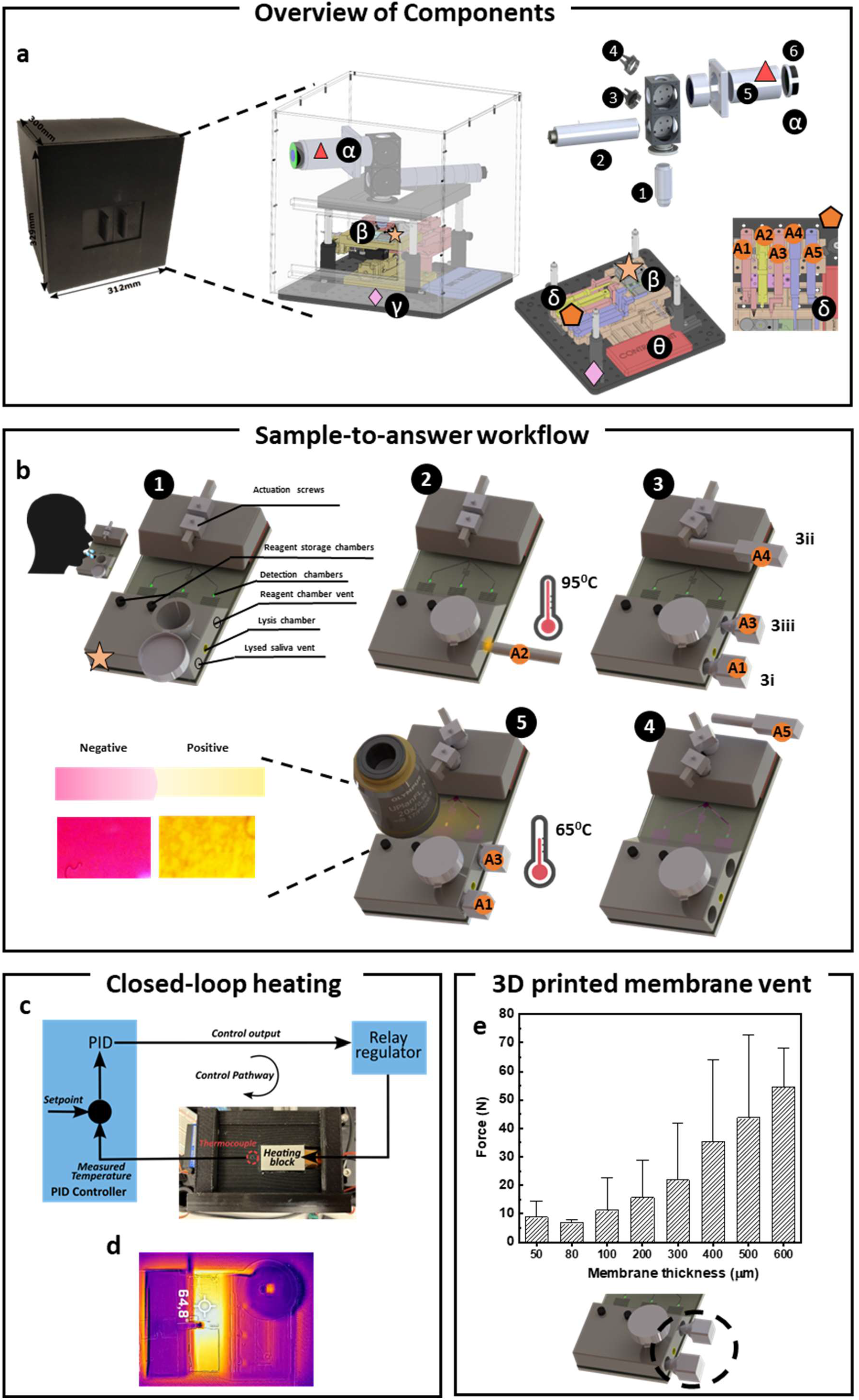
Assay automation. (a) The benchtop portable setup is depicted with all the internal components, color and letter coded with α-imaging module (PRICE), β-microfluidic cartridge, γ-the automation/control module. The exploded-view drawing of the off-the-shelf components of imaging module is shown in α. The components are number coded, (1) 20x objective, (2) illumination column, (3) beam splitter, (4) fully reflecting mirror, (5) tube lens, (6) CMOS sensor. The γ module further consists of δ-the linear actuator unit, and θ-the microcontroller and auxiliary components unit. The discreet components of the microfluidic cartridge (β) form the core of fluid handling and assay automation. (b) The linear actuator system (δ: A1-A5), heating units, and PRICE module sequentially interact with the microfluidic cartridge to enable a sample-to-answer workflow. Crucial components of automation, (c) Closed loop heating pathway enables heating to desired temperatures (d&e); (f) 3D printed micron-scale membrane for venting the lysis and reagent storage chambers are characterised in the range 50 µm-600 µm.

The operation of the microfluidic cartridge involves sequential steps (Fig. 2b), automated by a system of five linear actuators-δ (coded-A1, A2, A3, A4, A5), and heating modules controlled centrally by an Arduino UNO microcontroller (Fig. S7). As can be seen in Fig. 2b, following the collection of samples from the user, the saliva collection funnel is sealed with a lid (1). subsequently, the saliva is then heated to 95^0^ C for 3 min with heated tip fitted A2 to complete the lysis process (2). The temperature is controlled and monitored via cyclic linear movement of the actuator and infrared images, respectively. Following this, the vent membrane is ruptured with a sharp-tipped A1, to vent the lysis chamber (3i). A membrane thickness of 50 µm with a rupture force of 6.26 N (Fig. 2e) is chosen, owing to lower force, mechanical stability, and simple fabrication using the 3D printer. Next, the lysed sample is suctioned to the entrance of a Y-junction mixing channel, by releasing a compressed tunable elastomeric chamber triggered via actuator A4 (3ii). The reagent storage chambers are vented to atmospheric pressure via membrane rupture (3iii) with A3. The reagents are then mixed with lysed sample in a serpentine channel, via actuation (A5) of a second compressed elastomeric chamber (4). During the final step, the ruptured membranes are closed with A4 and A5 which are fitted with an O-ring to curb evaporation, and the cartridge is heated to 65^0^ C for amplification for 10 min (Fig. 2d). The temperature profile is achieved by employing a proportional–integral–derivative (PID) controller in conjunction with the ceramic heater (Fig. 2c). Subsequently, the colorimetric event is captured by PRICE centrally controlled by Raspberry Pi 4 microcontroller in tandem with Arduino microcontroller to scan the three detection chambers and different regions of each one. This movement is enabled by an X-Y translation stage directly controlled by the Arduino. For the X-translation, a CNC linear stage was employed. For the Y-translation a unique contraption involving a linear manual actuation stage and continuous servo motor. This concerted interplay between different modules is centrally controlled by the mobile application. Both the Raspberry Pi and Arduino execute commands via Bluetooth communication (Fig. S11). The end-point images demonstrate the presence of the target.

### 3.3. Imaging module development and characterization

LAMP technique was employed in conjugation with a nanostructured platform that plasmonically enhances the amplification reaction.^57^ In brief, the irradiation of ambient light on the plasmonic nanosurface causes a collective plasmonic oscillation of the free electrons (Fig. 3a). This results in non-radiative relaxation in the form of hot electron injection into the media, which has the potential to catalyze electron-driven nucleic acid amplification at the interface of the plasmonic surface and the media.^57^ In our previous work^57^, we demonstrated the acceleration of the amplification reaction by 9.15 times at a concentration of 1000 copies/µl, resulting in faster release of H^+^ ions hence changing the color of the pH dependent from fuchsia to yellow. Characterization of the color change of the pH-responsive amplification media over 60 min was demonstrated with a UV-Visible test, which was evident from the change of normalized reflectance (Fig. 3b).

**Fig 3.**
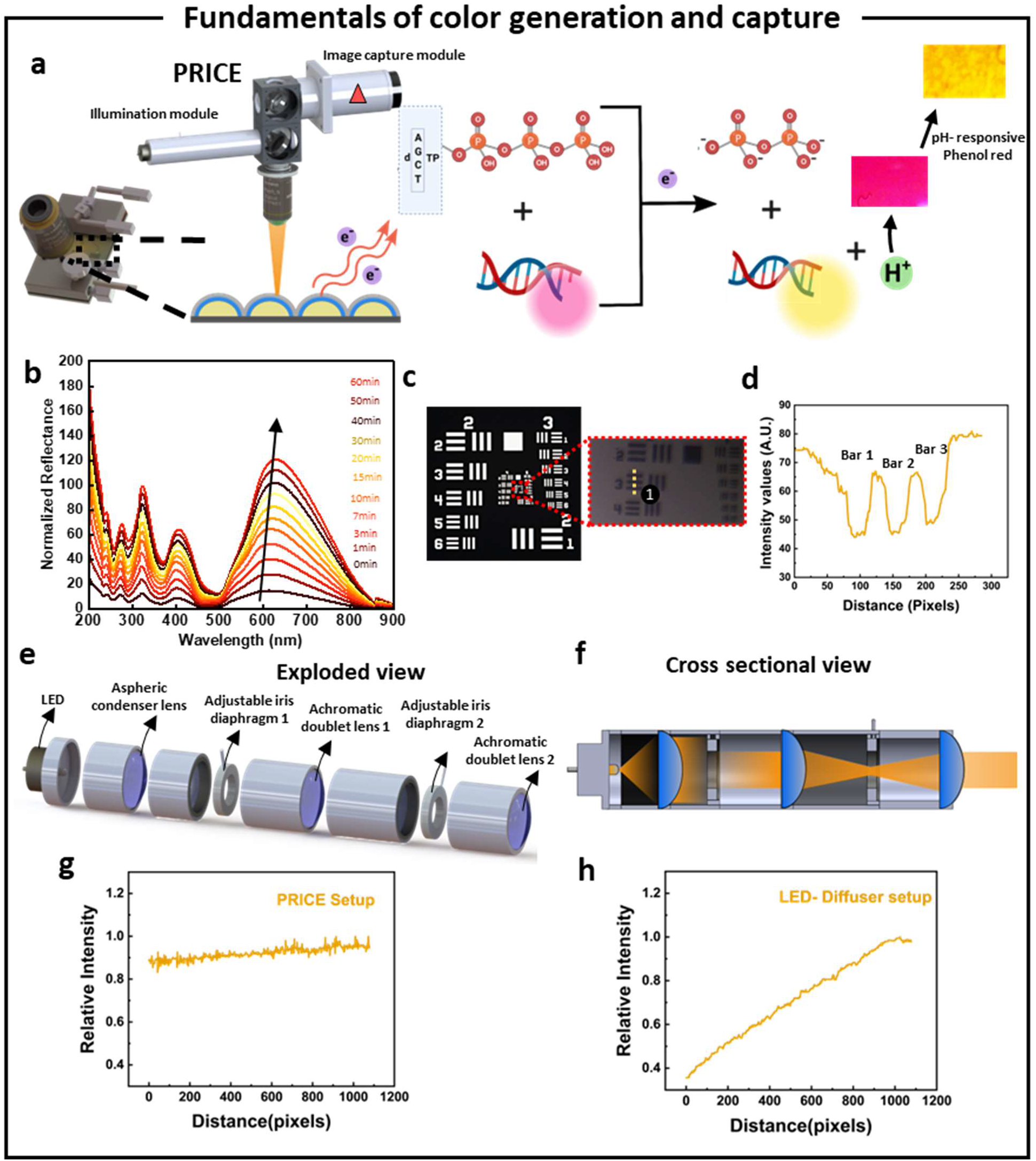
Optical module development and characterization. (a) The plasmonic nanostructured platform housed in the microfluidic chamber accelerates the pH dependent color change of the nucleic acid amplification assay upon irradiation of white light from the PRICE setup. (b) The UV-visible analysis of time-dependent color change of the amplification media for 60 min. (c) A UASF 1951 target imaged to characterize the FoV(d). Exploded view (e) and the light path (f) of the illumination column. The uniformity of irradiation of (g) PRICE outperformed the (h) LED-diffuser setup as offering better control over area and intensity of incident illumination.

This metallic nanosurface, which enhances the amplification reaction, is sensitive to illumination properties, which could ultimately affect the colorimetric readout. The PRICE module developed in this work provides uniform, consistent, and controlled illumination, comparable to a commercial brightfield microscope, while also being portable, cost-effective, and compatible with our automation setup. The PRICE module consists of two main submodules: (i) the illumination submodule, and (ii) the image capture submodule (Fig. 3a). The image capture submodule yielded a resolution of the 228 lp/mm or 4.4 µm using a 1951 US Air Force (USAF) resolution target (Fig. 3c, 3d). Subsequently, the FoV was also determined across the entire area of CMOS as 298 µm, enabling complete imaging of the detection chamber in 3 scans of different inter-chamber regions while not losing crucial information.

Source of illumination is an important property that can have predominant effect on the final color captured and observed.^63^ The main illumination design constraints include collimation, uniformity, spatial control, and numerical aperture (NA) of final incident light. To achieve these illumination characteristics, we drew inspiration from Koehler illumination setup (Fig. 3e, 3f), that is widely adopted in standard microscope illumination trains (Suppl. Info. N1).^64^ Moreover, Koehler illumination setup is more resilient to external disturbances like dust and optical imperfections, making them suitable for imaging with minimal variations over long and repeated cycles outside of controlled environments.^65^ The PRICE setup offered very uniform illumination where the relative intensity of the incident light varied only 17% (Fig. 3g, S6), whereas the regular LED-diffuser lens setup (LED placed in the focal plane of a diffuser lens) with an aspheric diffuser varied 65% (Fig. 3h), with a flat aluminium target.

### 3.4. Development and characterization of the microfluidic cartridge

Sample-to-answer workflow is realized by automating the sequential steps of the amplification assay--which include sample collection, lysis, mixing of reagents, and heating by employing our novel microfluidic cartridge. This multilayered microfluidic cartridge (Fig. 4a) leverages additive manufacturing to realise a sample to answer workflow. To obtain sequential metering of fluids in the microchannels, this work leverages elastomeric suction chambers that are tunable via a 3D printed screw-nut actuator contraption, obviating the need for expensive and bulky auxiliary equipment like syringe and peristaltic pumps (Fig. 4a). Mechanical actuation of these elastomeric suction chambers confers volumetric tunability, enabling flow manipulation in the microfluidic cartridge.

**Fig. 4.**
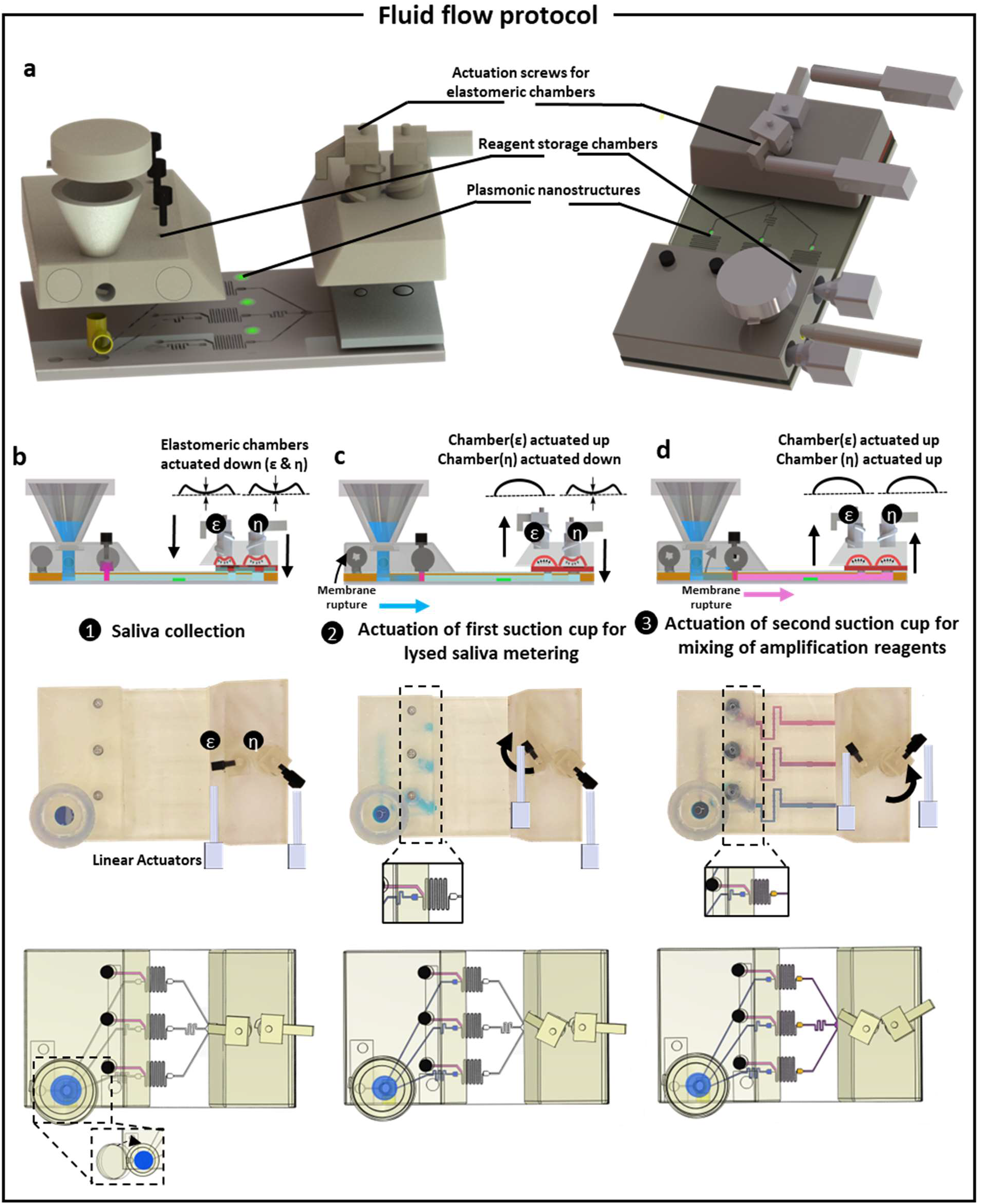
Fluid flow protocol. (a) Exploded view of the microfluidic cartridge. (b) Initially, amplification reagents were loaded into reagent storage compartments while the elastomeric chambers are in compressed state. Simultaneously, saliva was collected in a funnel for lysis and tightly sealed with a lid at the saliva inlet. (c) After completing the lysis step, the membrane was broken to vent the saliva channel, and the first elastomeric chamber was released from compression. (d) Once the saliva reached the Y-junction of the mixing channel, a second membrane was broken, exposing the reagent inlets to the atmosphere. The release of the second elastomeric chamber allowed the lysed saliva and reagents to enter the mixing channel.

The overall scheme and steps involved in the sample-to-answer operation as overviewed in Fig. 2b, is discussed in this section with emphasis around the microfluidic cartridge. The sequential fluid pumping steps required for amplification assay (Fig. 4b, 4c, 4d) is enabled by temporal actuation of the elastomeric suction chambers (steps b to d).All the steps (a to c) are facilitated via linear actuators housed inside the automated setup (Fig. 4a). In the first step (Fig. 4b), the sample is collected in a funnel while the microfluidic channels are under negative pressure as the elastomeric chambers are in compressed state. Subsequently, upon lysis by the heating tip fitted actuator, a thin 3D printed membrane is ruptured using sharp ended actuator to vent the lysed saliva chamber to atmospheric pressure. This venting enables the flow of lysed sample when the elastomeric chamber (ε) is actuated up (released state) leading to pressure-driven flow of the lysed sample to the mixing junction (Fig. 4c, inset showing the close-up view). In the final step (Fig. 4d), the second thin 3D printed membrane is ruptured to vent the reagent storage chambers to the atmospheric pressure, for mixing in the serpentine channels when the elastomeric chamber (η) is actuated up. The second membrane rupture vents the reagent storage enabling mixing with the reagents (Fig. 4d(inset)). The elastomeric chambers (Fig. 4(ε and η)) were designed in way that precisely match the volumes of the microchannels, in which the desired fluid flows upon actuation. The blue and red dyes were employed as analogue for saliva and reagents, respectively for the study.

The microchannels are designed and simulated using COMSOL to ensure mixing (Fig. 5a) and the ratio of lysed sample to reagent to be 1:10. Further, a uniform flow rate is ensured across the triplicate fluidic compartments for multiplexed detection assays (Fig. 5b). The total internal volume of the elastomeric chambers is a key design parameter as it affects the total volume of the fluid pumped. The elastomeric chambers work on the principle of air displacement generates a negative pressure in the microfluidic channels thereby inducing fluid flow along the pressure gradient.^41,65^ However, the mechanical contraption (Fig. S7) is prone to have residual volume under compressed state.^66^ Initially the chamber is assumed to be a hemisphere, hence the initial volume (V_in_) is given by^42^:

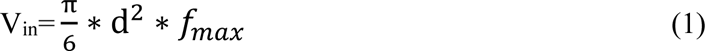

**Fig 5.**
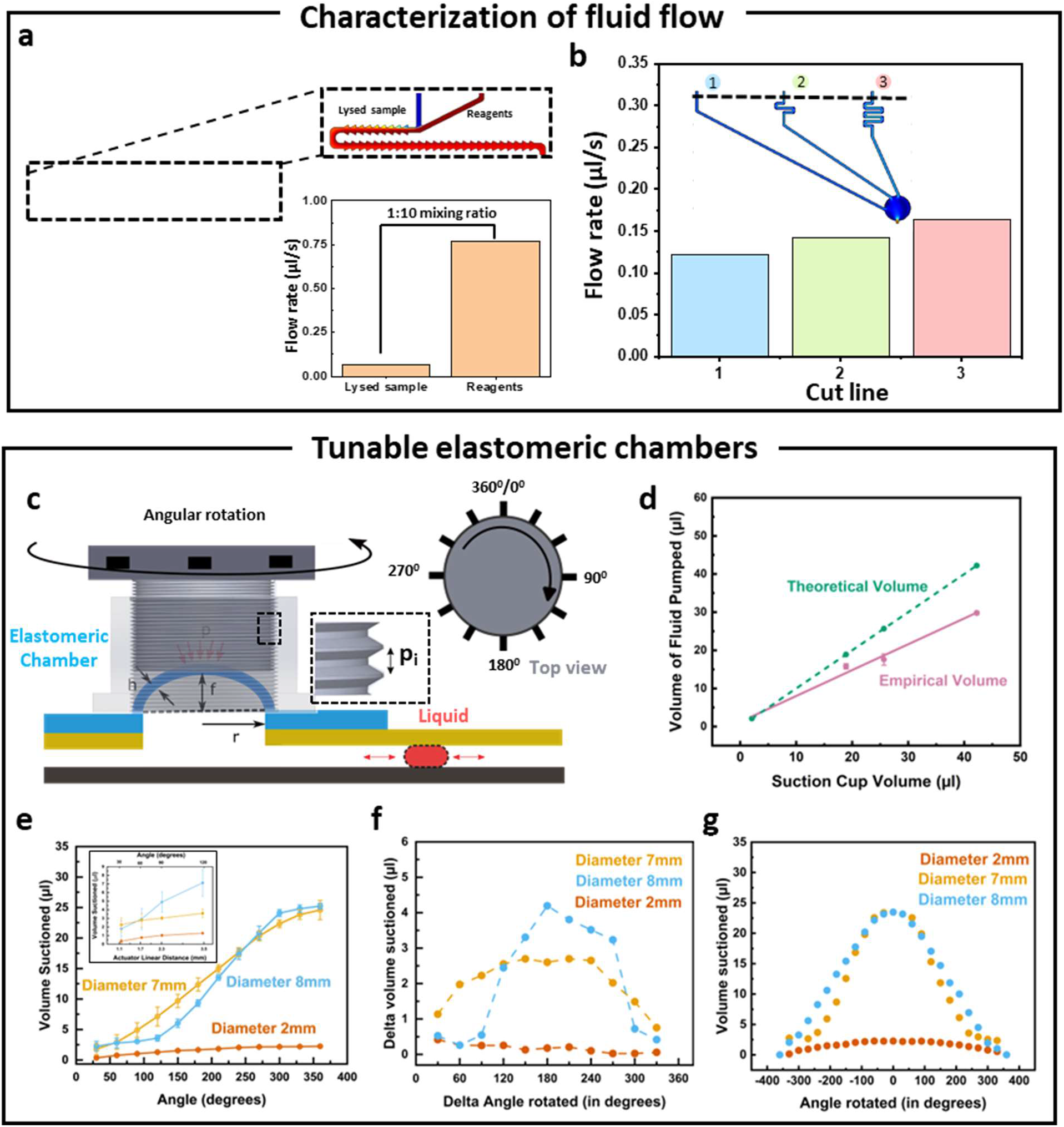
Microfluidic cartridge development and characterization. The microfluidic design parameters characterized with a COMSOL simulation studies for flow rate ratio of 1:10 (sample to reagents) (a) and pressure distribution (b). (c) The tunable elastomeric chambers operate using an additively screw-nut actuation setup, enabling control of fluid flow volume and direction through angular rotation from 0° to 360°. (d) Theoretical chamber volume is correlated with empirical volume pumped. (e) The graph illustrates the relationship between fluid volume pumped and screw angle for different elastomeric chamber sizes. Inset, depicts the correlation of volume suctioned to linear actuator distance moved. (f) Minimum manipulable fluid volume for a set screw rotation (360°) is characterized. (g) Repeatability studies demonstrate the precision of fluid volume pumped in one full pumping cycle (0° to 360°).

Here, d is the diameter of the elastomeric chamber and f_max_ is the central height of the hemisphere (Fig. 5c). The empirical volume of the fluid suctioned upon mechanical actuation deviates from the theoretical (actual) volume (Fig. 5d) pointing at the influence of residual volume upon complete actuation. Further, the empirical volume pumped varies linearly with the elastomeric chamber diameter (Fig. 5d). This enables fabrication of elastomeric chambers of desired volumes (accounted for residual volume) using simple and on-demand 3D printing.

To explore the precise tunability of the elastomeric chambers, screw-nut mechanical actuator (Fig. S6) was designed to manipulate the compression of the elastomeric chamber. For each rotation, the screw moves a distance equivalent to the pitch (Fig. 5c(P_i_)) which in turn manipulates the deflection of the elastomeric membrane (f) (Fig. 5c). This is demonstrated in Fig. S8, where the screw head is rotated in steps of 30^0^ either in clockwise (CW) or counterclockwise (CCW) direction to pump fluid in and out of the channels in multiple cycles. Upon testing with elastomeric chambers of different diameters (7 mm, 8 mm, and 2 mm), the trend between rotation angle and volume pumped (Fig. 5e) correlates with the analytical model, with a linear regime for f < h, and a non-linear trend for f>h (Suppl. Info. N2). Furthermore, the angle of the screw can be directly controlled via manipulating the linear distance moved by the linear actuator (Fig. S9). Hence the volume pumped is directly correlated with the linear distance moved by the actuator that can be easily controlled via a simple coding prompt (Fig. 5e, inset). This enables on-demand pumping of desired fluidic volumes and rapid design of elastomeric chambers at multitude of volumes and diameters. Capability of handling low fluidic volumes down to 0.1 µl per 30^0^ degree angular rotation was demonstrated with this mechanical contraption (Fig. 5f), showing precise tunability. The parabolic profile correlates to the analytical model (Suppl. Info. N1). Moreover, the system showed reversible bi-directional pumping capability (Fig. 5g), where a certain volume of liquid is pumped in and out of a microfluidic channel with great reliability and no volume loss in full pumping cycle (Fig. S8).

### 3.5. System validation

The clinical feasibility of the automated setup was demonstrated with a colorimetric LAMP assay for the detection of SARS-CoV-2 wild type (WT) RNA as a validation target. The colorimetric change of the amplification media is monitored via pH responsive dye, phenol red. In the event of amplification, the phenol red in the master mix undergoes a redox reaction mediated colorimetric change from fuchsia to yellow. The specificity of LAMP primers towards the SARS-CoV-2 wild type RNA is tested in Eppendorf tubes by incubating at 65^0^ C for 60 min. This is visually confirmed with a clear colorimetric change to vibrant yellow at a spiked SARS-CoV-2 WT RNA concentration of 8 ×10^5^ copies/µl (Fig. 6a).

Spiked SARS-CoV-2 WT RNA at different concentrations 8×10^5^ RNA copies/µl, 10^3^ RNA copies/µl, 10^2^ RNA copies/µl are loaded into the microfluidic cartridge and placed into the benchtop automated setup (Fig. 2a). The PRICE setup was employed to monitor colorimetric change at different time points (Fig. 6b). In brief, the CMOS sensor captures the image of the detection platform. The raw image is stored in the raspberry Pi for further processing. In addition to controlled illumination, the parameters of image capture, namely, white balance, gain (analog and digital) exposure time, framerate and ISO become important. The images are captured at a framerate of 30 frames per second (fps), an ISO of 295, and red and blue gains are set to 3 and 3 respectively. This color change from PRICE setup is visually juxtaposed with colorimetric change captured by a commercial brightfield microscope (Fig. 6b). The colorimetric signal, G^2^**/(**RxB)^57^, from both the setups were obtained and compared, where R, G, B are the average pixel values of the red, green, and blue channels, showing a shift in the signal at 10min. Subsequently, the colorimetric signal is relayed to a mobile phone application (Fig. S5). The PRICE setup offers a similar performance when compared with a commercial brightfield microscope (Fig. 6c). Clinically isolated RNA samples were tested with the platform (Fig. 6d). The colorimetric signal recorded shows statistically significant difference between healthy (S1, S2) and patient (S3, S4, S5) RNA isolates (Fig 6d & S10). The automated setup is operated via a mobile phone application which enables Bluetooth crosstalk between Arduino and Raspberry Pi. Further the user can track the real time status of the cartridge in the automated setup (Fig. S5).

**Fig 6.**
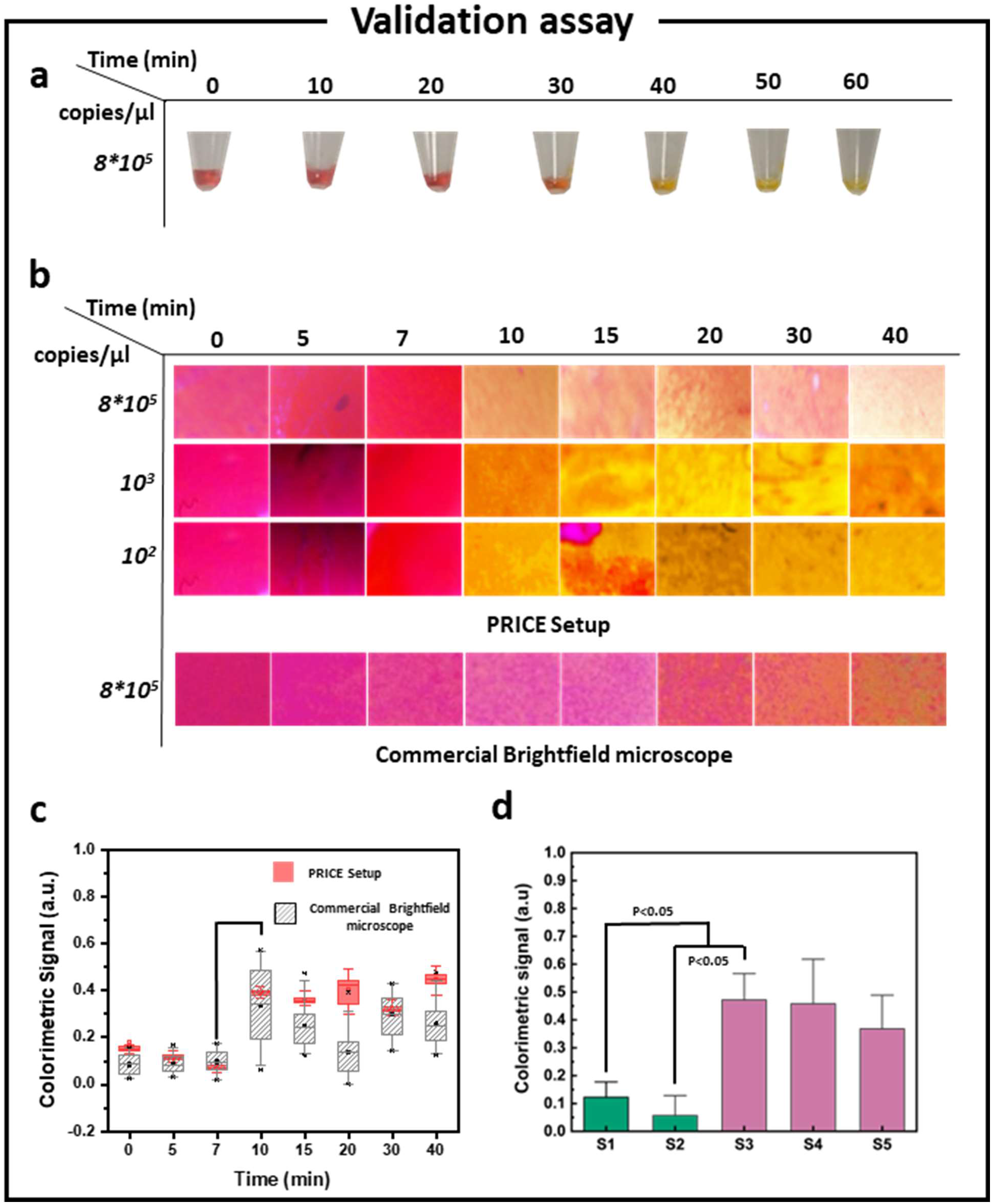
SARS-CoV-2 detection assay. The figure set represents a proof-of-concept study of a portable imaging setup for detecting SARS-CoV-2 at the point of care (POC) using phenol red-mediated colorimetric change. (a) Primers’ specificity is confirmed by conducting a 60-minute LAMP reaction in Eppendorf tubes. (b) Time dependent color matrix of various concentrations of RNA spiked. (c) The performance of the PRICE setup is compared with a commercial brightfield microscope. A graph illustrates the temporal change of the G^2^/R*B, referred to as the QolorEX signal, showing a significant change at 10 minutes. (d) Colorimetric signal plotted for different isolated RNA samples for healthy and patient samples.

## 4. Conclusion

In this work, we reported an automated setup for sample-to-answer colorimetric pathogen detection. The sequential steps of the amplification assay, signal transduction, and analysis were automated using three different modules, (i) an additively manufactured cartridge housing the novel angle-dependent tunable elastomeric chambers for precise microfluidic flow control, (ii) a portable reflected-light imaging setup with controlled epi-illumination (PRICE), and (iii) an automation/control and data analysis module. The specialized nanostructured platform recently developed in our previous work, that leverages plasmonic excitation for highly sensitive detection nucleic acid, was employed as the key detection technology. To capture the colorimetric change, PRICE was designed for imaging the assay chamber. The imaging setup offered superior spatial and spectral control with only a 17% variation in the relative intensity and a resolution and FoV of 4.4 µm and 298 µm, respectively. To eliminate the involvement of the user, the microfluidic cartridge, coupled with a set of actuators and mechanically actuated PDMS elastomeric chambers, was implemented by taking advantage of additive manufacturing techniques. The flow was shown to be mechanically actuated by a screw-nut mechanism with excellent control over the fluid pumping. This actuation mechanism demonstrated lowest volume of fluid suctioned at 0.1 µl for a 30^0^degree rotation of the actuating screw. The microfluidic chip also showed perfect extent in mixing lysed sample with the reagents. Regarding the final automation and control module, the cartridge operation was concerted using system of linear actuators and electrothermal heaters connected via Arduino UNO and Raspberry Pi microprocessors, which were remotely controlled via a smartphone application. The imaging was implemented in a direct comparison format with a Nikon brightfield microscope, and a colorimetric change was recorded in 13 min. In future iterations, the setup can be adopted to other detection and molecular assays that require sequential fluid handling steps. This automated system can truly enable the application of highly sensitive QolorEX platform at the POC for pathogen testing.

## Acknowledgments

The authors thank the Faculty of Engineering at McGill University, the Natural Science and Engineering Research Council of Canada (NSERC, G247765), the Canadian Institutes of Health Research (CIHR, 257352), the Canada Foundation for Innovation (CFI, G248924), and MI4 Emergency COVID-19 Research Funding (250611), for financial support. The authors acknowledge Nanotools-Microfab and the research facilities of NanoQAM at the Université du Québec à Montréal. SGY thanks Mitacs and McGill Engineering Doctoral Award for the scholarship funds.

## Author contributions

SM, and SGY contributed to the idea conception. SGY, and IIH contributed to the design, planning, and execution of the investigation. HS, AKJ contributed to execute the components of automation. TA, and MJ contributed to the fabrication and design of the plasmonic platforms. SGY, IIH, AKJ contributed to the preparation of the manuscript, with the support and collaboration of all co-authors. SM contributed to the project conception, design of the experiments, interpretation of results, resources and funding acquisition, and supervision of the work.

