## Supplementary Information for "Additive Manufacturing Leveraged Microfluidic Setup for Sample to Answer Colorimetric Detection of Pathogens"

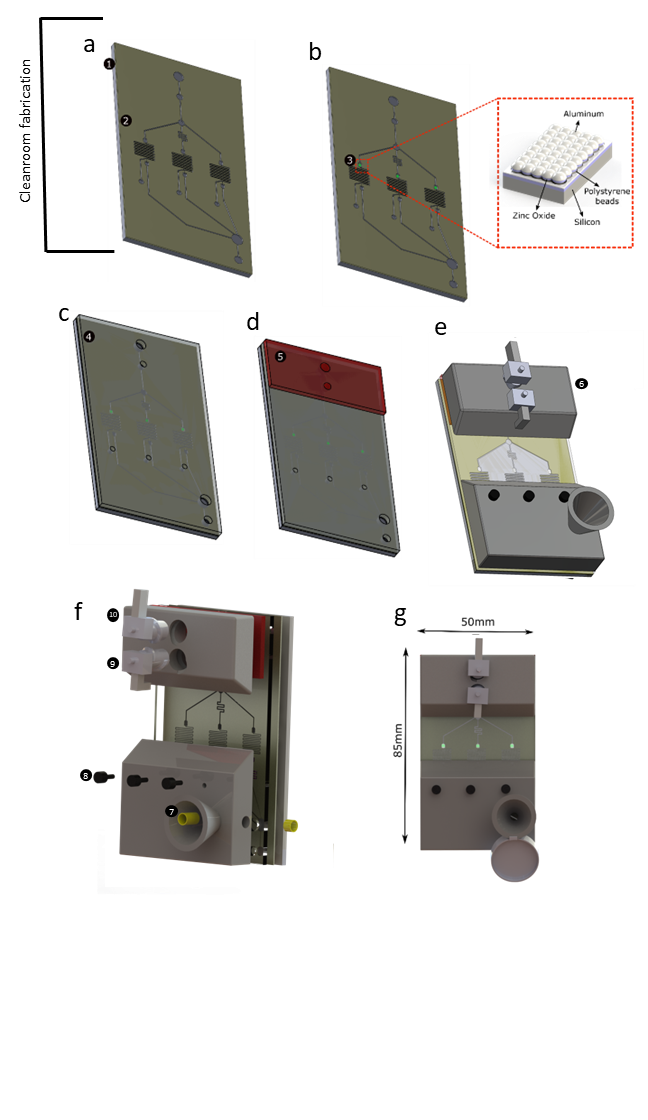

***Fig S1. Stepwise fabrication process*** (a) The first step of the fabrication process is the patterning of the fluidic channels using UV photolithography process. A 50micron thick SU-8 layer (2) was patterned on a silicon substrate (1) via a straightforward lithography process. (b) In the second step, the color sensitive platform (3) was fabricated using a fabless nano-patterning technique. The subsequent fabrication processes are performed out the cleanroom. (c) A thin PDMS layer (4) is bonded to the SU-8 fluidic layer via plasma bonding to created closed microfluidic channels with punched holes at channel inlets. (d) Next, PDMS based elastomeric chambers (5) fabricated using SLA printed molds are bonded to the fluidic-PDMS layer by plasma bonding. (e&f) Finally, the SLA 3D printed cartridge (6) is bonded to the PDMS covered microfluidic chip using a double-sided tape conducive to plasma activated bonding. The figure (f) shows the exploded view of the cartridge showing multiple components. The brass metal inserts (7) are employed for lysis of the sample. Small screws (8) are used to seal-off the reagent storage chambers. The components-9&10- are 3D printed screws to holding the elastomeric chambers in place and subsequently actuated for flow control in the cartridge.

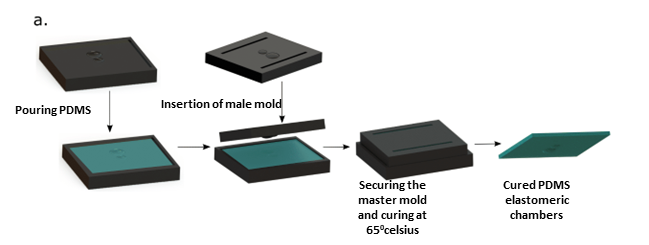

***Fig S2. Elastomeric chamber fabrication protocol.*** (a) The elastomeric chambers are fabricated using SLA printed molds. The 3D printed molds are first surface treated to avoid curing inhibition at the PDMS mold interface. The high resolution of the printing ensured low roughness of the PDMS thereby ensuring strong bonding.

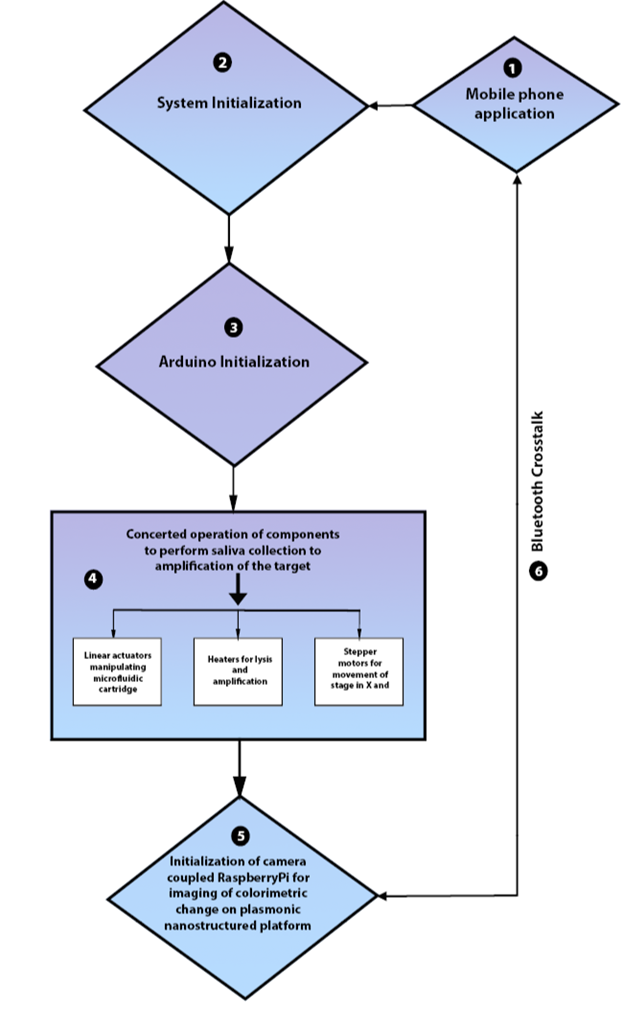

***Fig S3. Overview of high-level system design*** (a). Flow diagram depicting the high-level operation of microfluidic sample-to-answer pathogen detection platform, with major emphasis on electronic components involved. The number coded blocks indicate the sequence of the individual process involved in the operation. In brief, Step-1 to 3 involves the user to connect to the system via a mobile application enabled by a on-board Bluetooth module connected to the Arduino UNO. In the next subsequent step, the components in the Arduino are activated in concerted sub steps to complete the processes from sample collection to LAMP reaction incubation. Following the completion of the reaction, the Bluetooth communicates with Raspberry Pi via Bluetooth to initiate imaging and subsequent movement of the stage for multiplexed pathogen detection.

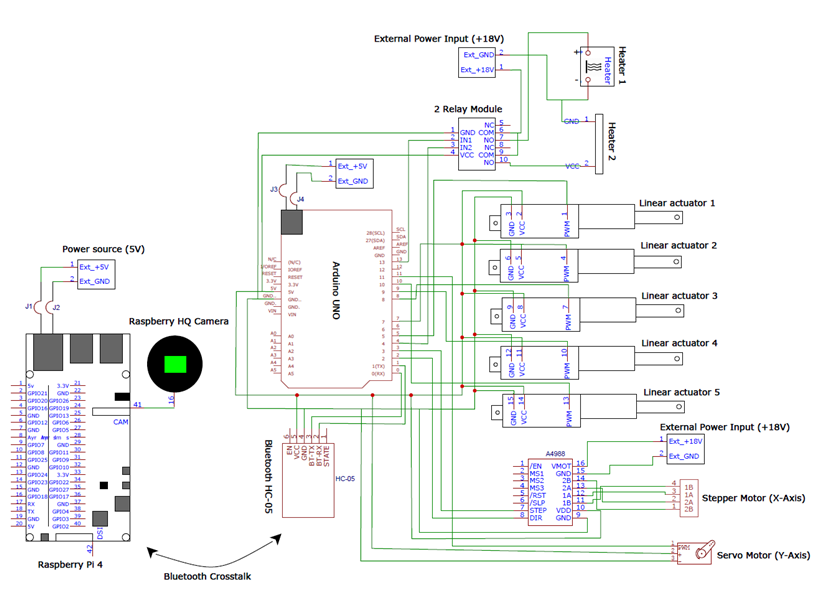

***Fig S4. Overview of electronics.*** The figure shows the electronic CAD layout of all the electronic components enabling the automation of the sample to answer processing of the system. The major constituents involved in the prototype controlled by Arduino UNO and Raspberry Pi are, (i). actuator system (five in number to automate the process of saliva collection and sequential delivery of reagents typically involved in a LAMP assay), (ii). Two DC heating components to controlled heating for pathogen lysis and subsequent nucleic acid amplification reaction, (iii). Raspberry Pi 4 (4GB RAM) coupled with CMOS sensor for imaging of the colorimetric change, (iv). A HC-05 Bluetooth module that enables cross talk between the mobile application and the Arduino UNO, and subsequently between Arduino UNO and Raspberry Pi 4.

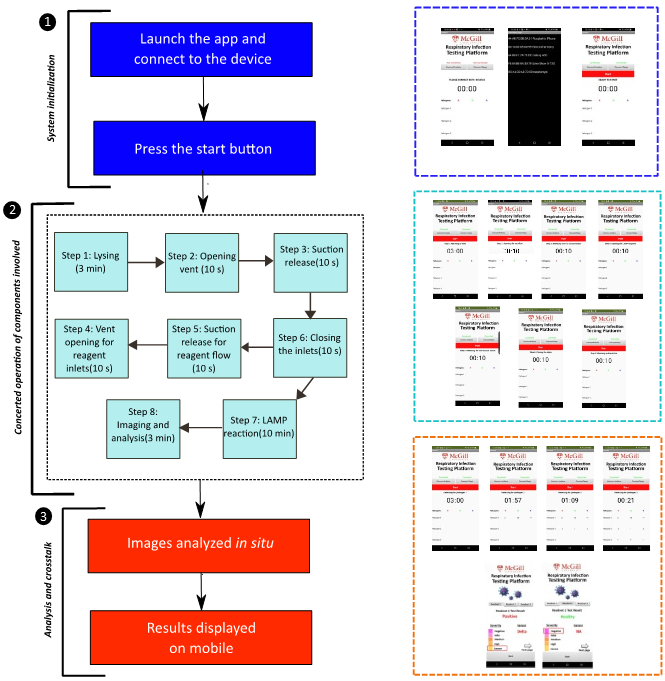

***Fig S5. Overview of operation of the mobile enabled automation***. This figure gives an overview of process flow involved in the operation of the mobile application working in tandem with two major microcontroller modules, namely, Arduino UNO and Raspberry Pi 4. The operation of the application can be broadly categorized into three main subsets, (1) System initialization- The crosstalk between the mobile phone and the microcontroller modules is enabled by a stable Bluetooth connection. Hence the first step is establishing a connection between mobile and the modules and subsequently press the button labelled ‘Start’ to send a message to the Arduino and Raspberry Pi 4 to start initialize the process (color coded in navy blue). Following this, in subset (2), the components start operating in a concerted fashion. Each component substep is are clocked to enable operation in the defined sequence. The user can monitor the progression of the process on the mobile application (color coded in light blue). Once the assay incubation time is completed, the images are captured and analyzed in substep (3). In brief, RGB values are extracted from the images and analyzed further to display the final result on the screen easily interpretable by the user (color coded in red).

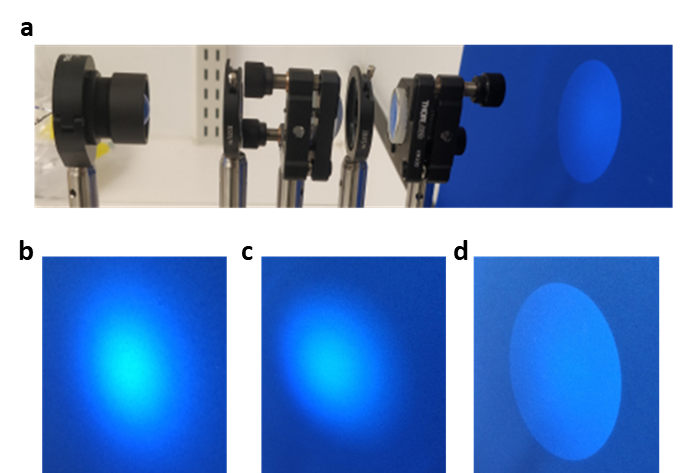

***Fig S6. Components of illumination*** (a) The benchtop setup of the illumination column for visual characterization. The collimated light beam is irradiated onto a blue background and subsequently images were captured with phone camera. The visual intensity profile of the collimated beam is shown for, (b) simple LED with no lens setup, (c) LED-diffusing lens setup, (d) PRICE illumination column.

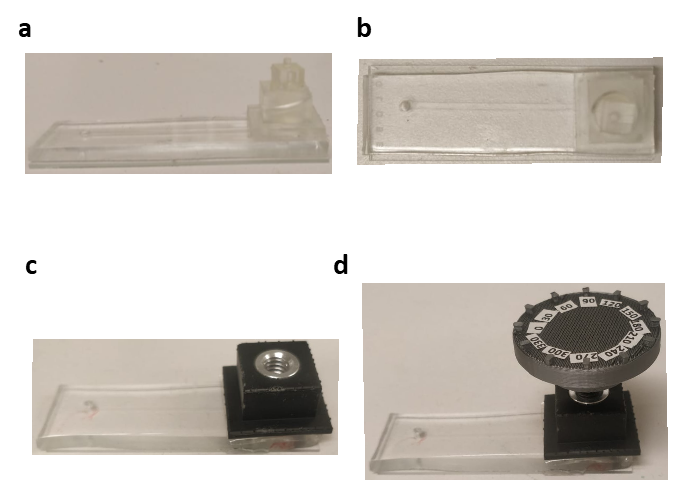

***Fig S7. Depiction of experimental setup for suction cup characterization***. (a&b) 3D printed module employed for characterization of empirical volume of the suction cups. (c&d) The fluidics setup employed for demonstration of fluid manipulation.

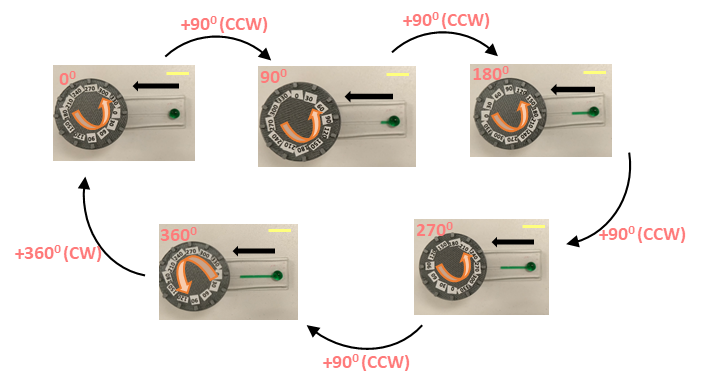

***Fig S8. Demonstration of various of aspects of fluid flow manipulation***. (i) The fluid pumping is depicted by the controlled mechanical actuation of the PDMS suction cup. The tip of the screw exerts a perpendicular force on the suction cup displacing/releasing the air underneath causing a pressure gradient conducive to fluid flow. The demonstration shows volumetric fluid metering based on the angle of the screw rotated in either CW or CCW directions (Scale bar- 15mm).

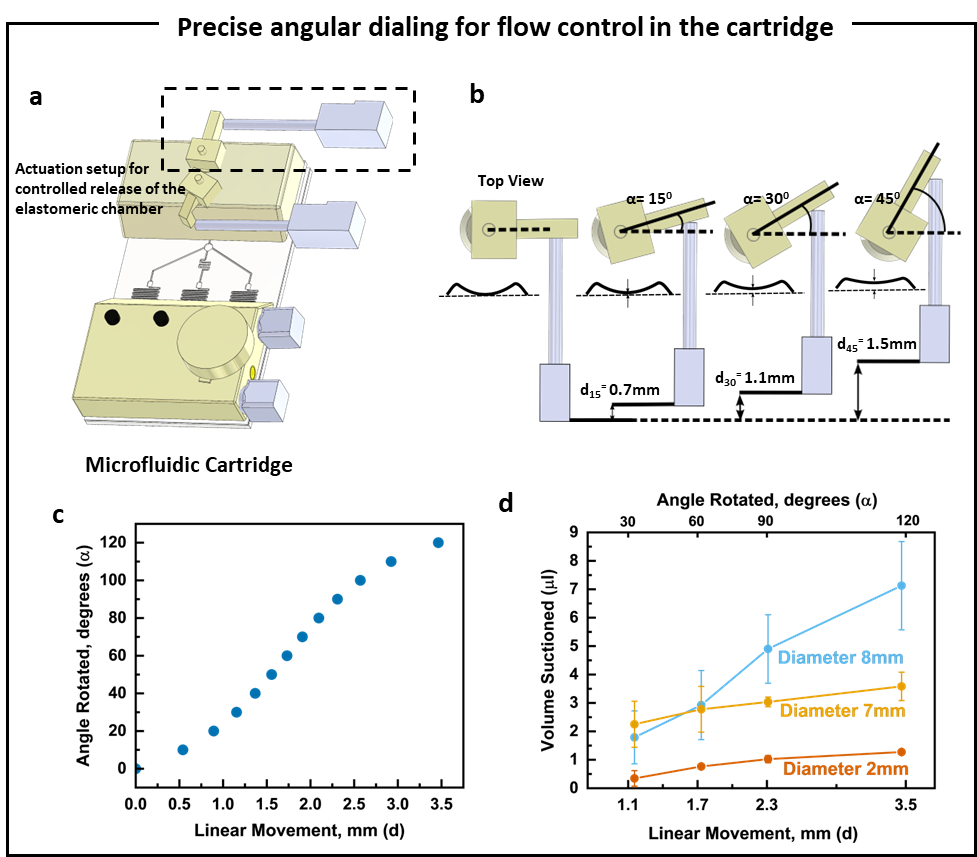

***Fig S9. Precise angular dialing demonstration.*** (a) The control and data analysis module houses a set of linear actuators that interact with the microfluidic cartridge. (b) The actuation setup for the elastomeric chambers involves a linear actuator interacting with a rotating screw. (c) The angular rotation is precisely controlled via linear movement of the actuator, following a set trend (d).

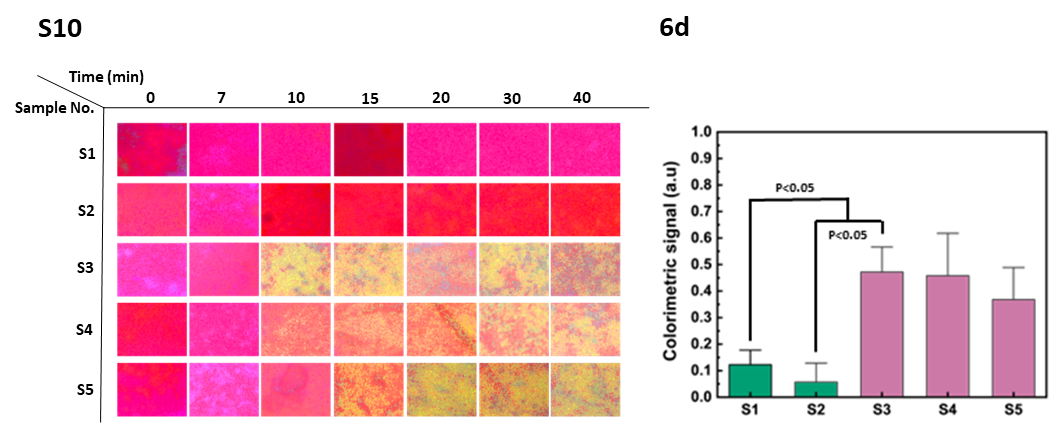

***Fig S10.*** Image matrix depicting time-dependent colorimetric change for RNA isolates (S1 to S5).

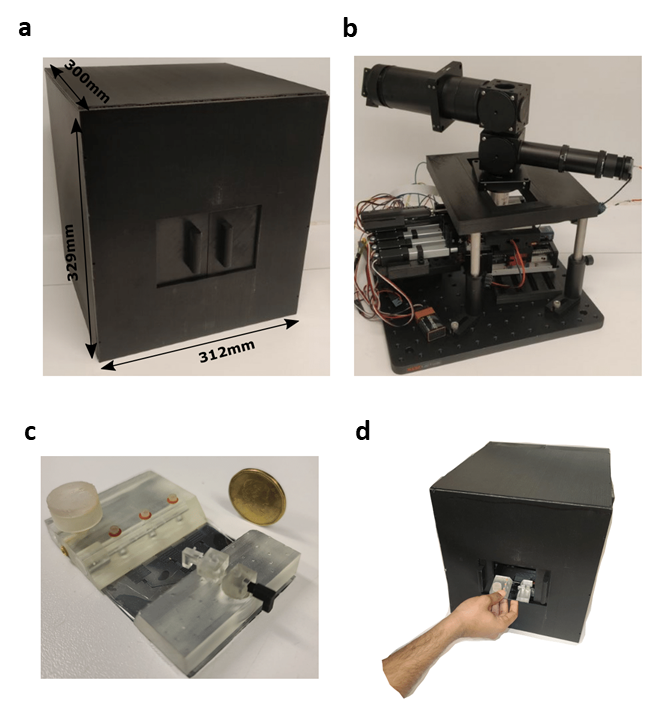

***Fig S11. Camera captured pictures of the prototype*** (a). Picture showing the isometric view of the outer enclosure of the setup with dimensions specified. (b)-(d). Different views of the inner assembly and wiring of the setup, (b)- Front view, (c)- Isometric view and (d)- Top view.

**Note-N1**

In the proposed design for the illumination submodule, the NA of the incident light on the sensing chamber is dictated by the NA of the 20x objective when illuminated with collimated beam. In brief, a 5000 K 90CRI LED (total viewing angle of 60^0^) is placed at the focal plane of an aspheric condenser lens with a diffuser (Fig.S6, Fig.3e). The exiting beam illuminates an adjustable iris diaphragm-1 allowing control over the area of illumination in relation to radius of diaphragm-1 opening. The partially collimated beam is then focused on an achromatic doublet lens-1 that forms real image of the light source on a diaphragm-2. The diaphragm-2 opening dictates control over the intensity of illumination. In the final step, the real image comes out as a collimated beam of light after passing through another achromatic lens-2 placed at the focal length.

**Note- N2**

We hypothesized that the volume of fluid suctioned/pumped (V­_p_) can be varied by manipulating the deflection of the PDMS membrane (f) (Fig.4c). This volume, V_p_, is controlled by the mechanical actuation of a screw and nut system (Fig.S7). The central idea revolves around the concept that the volume, V_p_, can be controlled via fine threaded screws mechanically pressing on the buttons. For each rotation, the screw moves a distance equivalent to the pitch which in turn manipulates the deflection of the suction membrane (f). This is demonstrated in Fig.S8, where the screw head is rotated clockwise (CW) and counterclockwise (CCW) to pump fluid in and out of the channels in multiple cycles. The volume of the spherical suction cup is given by the equation^1^,

$V= \frac{\pi f}{6}(3r^{2}+f^{2}$)	(1)

Where V is the volume stored by the suction cup, f is the deflection, r is the radius of the suction cup. The deflection f is in turn a function of applied pressure and for silicone polymer (Poisson’s ratio of 0.5) is given by,

$f=\left( \frac{9r^{4}}{64Eh^{3}} \right)p, for f\leq h$	(2)

$f=\left( \frac{3r^{4}}{16Eh^{4}} \right)^{1/3}hp^{1/3}, for f>h$	(3)

Here E is the elastic modulus of the chamber material, h is the chamber PDMS membrane thickness, and p is the loading pressure. These equations hold true only when 2r>h, which is the case in our current work. From equations, (1), (2) and (3), it can be inferred that volume of the suction cup (v) varies cubically with loading pressure (p) in the f ≤ h and varies non-linearly in the regime f > h. The experimental results correlate with the theoretical profile as in Fig.4d. Specifically, we demonstrated the control of fluid flow with actuation varying in steps of 30^0^ angle either CW or CCW. Different sizes of suction cups (Diameters (d)- 2mm, 7mm, 8mm and height(h)- 1mm) were employed to demonstrate this controlled flow actuation, Fig.4. Given the pitch of the screw used is 1.4mm (distance moved by the screw in 1 full rotation), the volume suctioned plateaus after 300^0^ of rotation.

The delta volume suctioned is given by first order differential of the equation (4), given by,

$\frac{dV}{df}= \frac{\pi}{2}(r^{2}+f^{2})$	(4)

This equation shows that change in suction cup volume (i.e volume of fluid suctioned), is in quadratic relationship with the deflection. The experimental results are in correlation with this as demonstrated in Fig.4f, showing a parabolic profile. In other words, the angular control allows for precise manipulation of the fluid volume in the channel. By incorporating this screw-nut actuation system with a screw of pitch 1.4mm, we demonstrated pump of fluid volumes as low as 0.1 µl (Fig.4f). This process of fluid pumping into the channel is found out to be in correlation with the fluid pumping out of the channel in one full cycle (Fig.4g). This precise and reliable manipulation of fluid could have potential implications for facile metering of fluid at very low volumes with minimal user training.

**Note- N3**

Prior studies^2,3^ have extensively reported strategies to extend the shelf-life of the LAMP reagents under different storage conditions. For example, Kaymaz et al.^2^ reported a shelf life of 3 days at room temperature and 6 days at 4^0^C, when the *Bst* polymerase and nucleic acid sample are excluded. Lee et al.^3^ reported loss of the amplification activity in one day, whereas the reagent shelf life went up to 45 days with the addition of sucrose for polymerase stabilization. Similar strategies were reported using lyophilization and drying with addition of stabilizing compounds like trehalose^4^.
